# CircRNA-Pro: A Novel Toolkit for High-Precision Detection of Differentially Expressed Circular RNAs and Translatable Circular RNAs

**DOI:** 10.1101/2024.03.13.584785

**Authors:** Wei Song, Liqun Yu, Tianrui Ye, Honglei Zhang, Yan Wang, Yang Yang, Dawei Shen, Weilan Piao, Hua Jin

## Abstract

With the increasing discovery of circular RNAs (circRNAs) and their critical roles in gene regulation and disease progression, there is a growing need for more accurate and efficient tools for circRNAs research. In response, we have developed an integrated software suite specifically for circRNAs. This all-in-one tool specializes in detecting differentially expressed circRNAs, including those with the potential to be translated into proteins, and allows for comparing against relevant databases, thereby enabling comprehensive circRNA profiling and annotation. To enhance the accuracy in detecting differentially expressed circRNAs, we incorporated three different software algorithms and cross-validated their results through mutual verification. Additionally, this toolkit improves the effectiveness in identifying translatable circRNAs by optimizing Ribo-seq alignment and verifying against public circRNA databases. The performance of circRNA-pro has been evaluated through its application to public RNA-seq and Ribo-seq datasets on breast cancer and SARS-CoV-2 infected cells, and the results obtained have been validated against previous literature and databases. Overall, our integrated toolkit provides a reliable workflow for circRNA research, facilitating insights into their diverse roles across life sciences.

## 1. Introduction

Circular RNAs (circRNAs), distinct from traditional linear RNAs, are characterized by their unique covalently closed loop structure [1-3]. This structure contributes to their increased stability, which is essential for their significant roles in various cellular processes [4-8]. CircRNAs are involved in gene regulation, playing roles in the development of diseases such as cancer [9-11], neurodegenerative disorders [12-15], and are crucial for normal cellular development [16-18]. Recent studies have shown their roles as microRNA sponges [19, 20], RNA-binding protein sponges [21, 22], and nuclear transcriptional regulators [23], highlighting their diverse functions in cellular biology. Furthermore, emerging research has revealed that a subset of circRNAs possesses the capacity for translation [24-27], uncovering novel biological functions and deepening our understanding of their diverse roles in cellular processes.

Despite recent advances in the detection of circular RNAs [28-30], several challenges remain to be addressed, including inconsistencies in results due to differences in algorithms and the need to improve the accuracy of detection results [31-33]. The development of more precise and efficient software is crucial for a deeper understanding of the biological functions of circRNAs. Currently, there are numerous software tools available for the detection of circRNAs (e.g., CIRI2[34], CIRCexplorer2 [35], CirComPara2 [36]). However, there is a limited number of tools specifically designed for the detection of differentially expressed circRNAs, with CIRIquant [37], DEBKS [38], and Circtest [39] being the primary options. Due to the utilization of different algorithms, these tools often produce inconsistent results [32, 38], making it more challenging to accurately identify differentially expressed circRNAs. Currently, the most widely used method for detecting translatable circRNAs is to align the Ribo-seq sequencing reads with the sequences flanking the circRNA splicing junction sites to identify them [40]. However, these methods overlook an important issue when detecting translatable circRNAs: they fail to remove the sequences flanking the circRNA splicing junctions that can be mapped to the genome before performing the alignment with Ribo-seq reads [41, 42]. This can lead to inaccurate determination of whether the Ribo-seq reads mapped to these regions originate from circRNAs or the genome. Meanwhile, several databases related to translatable circRNAs have emerged, and integrating the conventional Ribo-seq alignment method with database verification strategy may enhance the accuracy of detection results.

To address the existing limitations in circRNA research tools, we have developed circRNA-pro, a comprehensive and user-friendly software package. This tool can not only predict circRNAs but also detect differentially expressed circRNAs, identifiy potentially translatable circRNAs, and allows for comparing against relevant circRNAs databases. To enhance the accuracy of detecting differentially expressed circRNAs, circRNA-pro integrates three distinct software algorithms, with cross-validation of their results. Additionally, it optimizes the alignment of Ribo-seq reads with circRNA sequences and verifies the results against well-established translatable circRNA databases to improve the accuracy of predicting translatable circRNAs. To facilitate use by researchers with limited bioinformatics expertise, circRNA-pro is packaged into a Docker image, allowing for easy distribution and execution across various computing platforms, greatly facilitating its use by researchers and potentially increasing the efficiency of circRNAs research. The performance of circRNA-pro was evaluated using publicly available RNA-seq and Ribo-seq data from breast cancer cells and SARS-CoV-2 infected cells. The test results demonstrated that circRNA-pro can accurately detect differentially expressed circRNAs as well as translatable circRNAs, with subsequent validation found in the literature and relevant circRNA databases. In summary, the development of CircRNA-pro will significantly simplify and accelerate circRNA research by providing a user-friendly and accurate tool for the detection and analysis of differentially expressed and potentially translatable circRNAs.

## 2. Results

### 2.1 Overview of circRNA-pro

#### The detection of all circular RNAs

The detection of circRNAs from sequencing samples begins with the quality assessment of the total RNA sequencing data using FastQC. FastQC enables a rapid overview of the data quality including read quality, GC content and other metrics from high-throughput sequencing. The raw reads are then processed with Fastp for quality control, including adapter trimming and filtering of low-quality bases. The cleaned reads were aligned to the reference genome using BWA-MEM [43], with the reference genome downloaded from GENCODE. Aligned reads are subsequently analyzed with the CIRI2 [34] to identify circRNAs across various samples by detecting reverse-mapped junction reads that are aligned to genomes in a non-collinear order (Fig. 1).

**Fig. 1.**
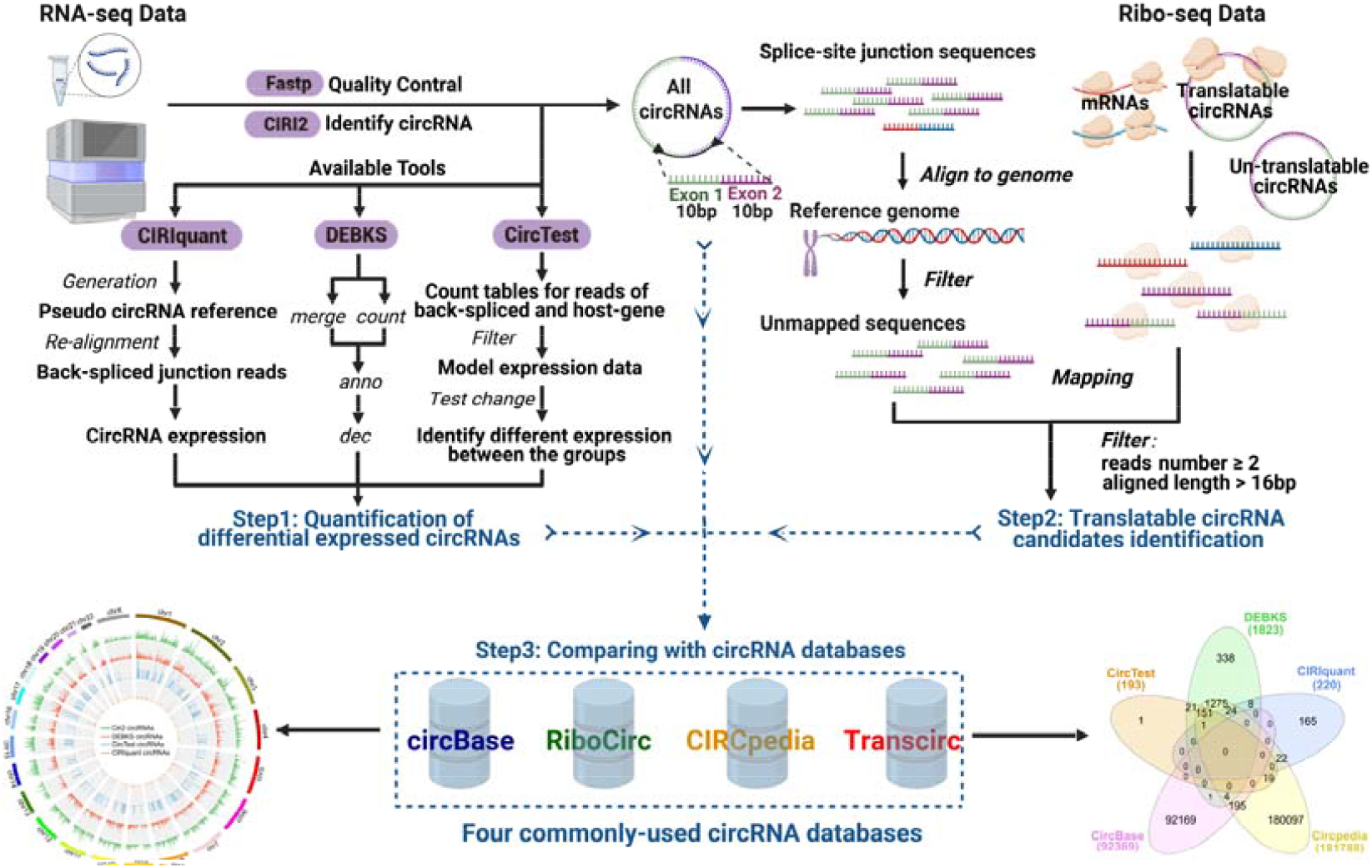
The workflow of circRNA-pro for detecting differentially expressed circRNAs and translatable circRNAs with total RNA-seq and Ribo-seq data.

#### The detection of differentially expressed circular RNAs

To identify differentially expressed circRNAs, we adopted a multi-algorithm strategy by utilizing three distinct programs: CIRIquant, DEBKS and CircTest, as demonstrated in step 1 of Figure 1. CIRIquant quantifies circRNAs expression by calculating the ratio of back-splicing junction reads to host gene reads [37]. On the other hand, DEBKS first detects back-splicing events, then performs differential expression analysis on these back-splicing events using edgeR [38]. Since edgeR is incompatible with direct use of back-splicing ratios, DEBKS reports differentially expressed back-splicing events rather than circRNAs [38]. CircTest represents circRNAs expression as the proportion of back-splicing junction reads out of all measurable junction reads, and performs statistical testing to identify differential circRNAs expression [39]. By integrating the results from these different tools, the reliability of the differentially expressed circRNAs detection results can be enhanced through cross-validation between various software.

#### The detection of translatable circular RNAs

To detect translatable circular RNAs, we employed a combined approach that involves aligning Ribo-seq reads with the sequences flanking circRNA splice sites and comparing the identified circRNAs with well-established databases of known translatable circRNAs (step 2, Fig. 1). Initially, we extracted 10 bp sequences flanking the circRNAs splice sites and concatenated these to create junction-spanning sequences. The junction-spanning sequences were then aligned to the reference genome, and sequences that could be aligned to genome were filtered out. These non-alignable sequences represented the unique splice junction repertoires of the circRNAs. Subsequently, Ribo-seq reads that could not be aligned to the reference genome were aligned to the junction-spanning sequences of the circRNAs. CircRNAs with two or more Ribo-seq reads mapped were considered as potentially translatable circRNAs candidates. These potentially translatable circRNAs candidates will be compared with two commonly-used databases of translatable circRNAs in the next step to confirm the detection accuracy.

#### The comparison of detected circRNAs with public circRNAs databases

To comprehensively assess and validate the circRNAs detected by the circRNA-pro software, we compared these circRNAs against four widely-used circRNAs databases: CircBase, Circpedia, Ribocirc, and TransCirc (step 3, Fig. 1). CircBase and Circpedia encompass a diverse range of circRNAs identified in multiple species, while Ribocirc and TransCirc are more specialized, focusing specifically on circRNAs that exhibit translational capabilities. Through comparison with these databases, we will further validate the identified translatable circRNAs candidates, as well as distinguish the known and novel circRNAs.

### 2.2 The comparison of four well-known circRNAs database

CircBase, Circpedia, RiboCirc, and TransCirc are four key databases critical to research on circular RNAs. Despite their importance, a comprehensive comparative analysis across these major circRNAs databases remains scarce. To address this gap, we conducted a comparison focusing on human circRNAs as cataloged in these databases. We considered circRNAs with genomic coordinate discrepancies of less than 10 bp across the databases to be identical. The comparison results showed that TransCirc contains the largest number of circRNAs at 328,080, while RiboCirc has the smallest collection at 1,201 circRNAs (Fig. 2A). For most circRNAs in the databases, the genomic distances between the back-splice sites are between 1000-50000bp (Fig. 2B). 80% of the exonic circRNAs have lengths less than 2500 bp (Fig. 2C). Most of the circRNAs in the four databases contain 1-2 genes (Fig. 2D). Approximately 70% of circRNAs in the four databases contain 1-10 exons (Fig. 2E).

**Fig. 2.**
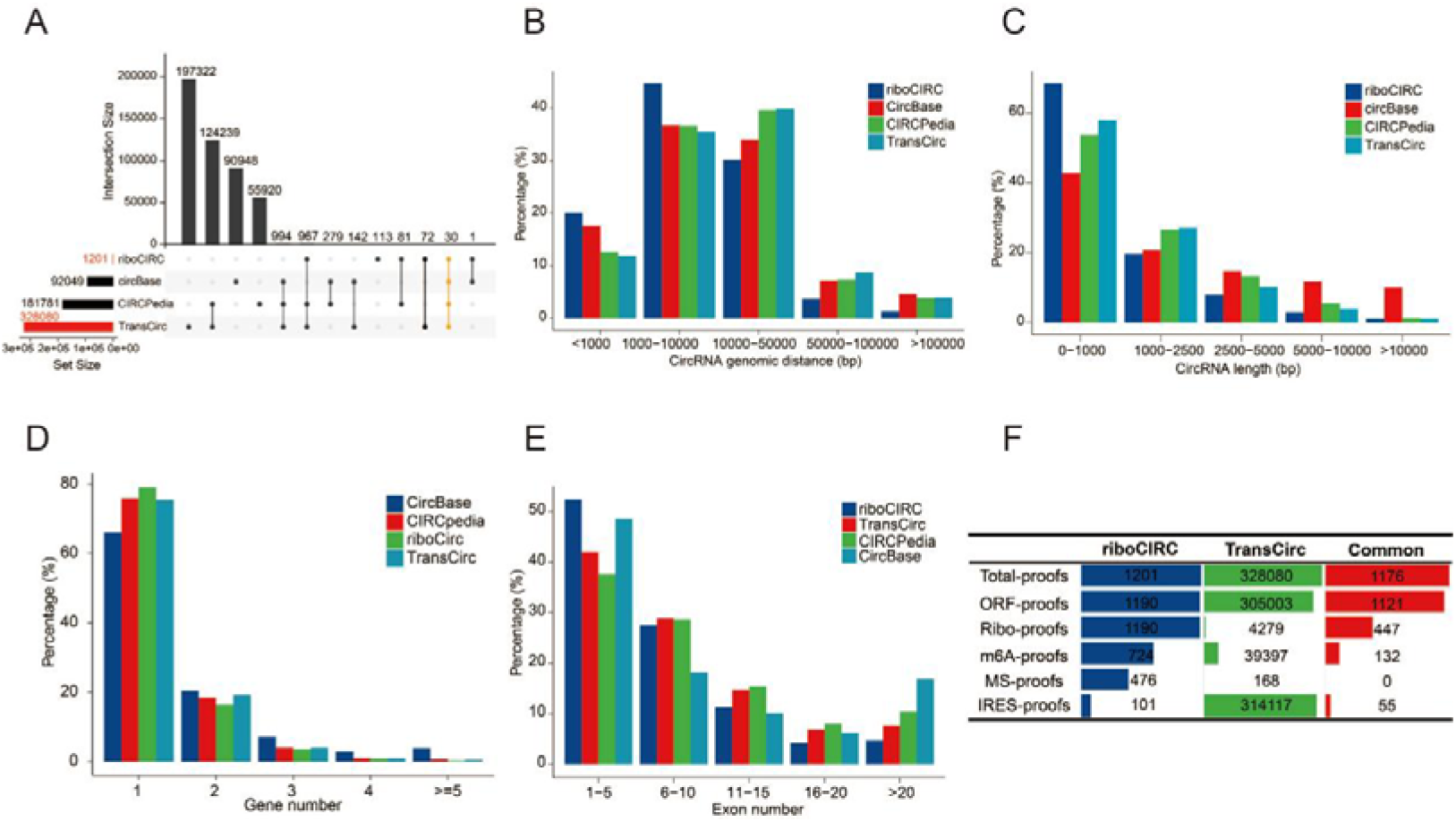
Comparison of circRNAs across different circRNA databases. (A) Intersection of circRNAs across different databases. (B) The genomic distance between back-splice sites of circRNAs. (C) The distribution of exonic circRNAs length. (D) Average number of genes per circRNA. (E) Average number of exons per circRNA. (F) Comparison of circRNAs with varying support evidence in RiboCIRC and TransCirc.

RiboCirc and TransCirc are two well-known databases containing circRNAs with potential translational capabilities. Each circRNAs entry in these databases is supported by various support evidence, such as Open Reading Frames (ORFs), m6A methylation modifications, Mass Spectrometry (MS) data, Internal Ribosome Entry Sites (IRES), and Ribo-seq data. To better understand the differences between these databases, we compared the circRNAs and their supporting evidence in RiboCirc and TransCirc (Figure 2F). The number of circRNAs in TransCirc significantly exceeds that of RiboCirc. CircRNAs supported by ORF evidence constitute the majority in both databases. Additionally, RiboCirc contains 476 circRNAs that are supported by both Ribo-seq and MS evidence, while TransCirc includes 3 circRNAs with both Ribo-seq and MS evidence.

### 2.3 The performance of circRNA-pro on the data of breast cancer

To assess the performance of circRNA-pro, we evaluated its accuracy on publicly available RNA-seq and Ribo-seq data derived from breast cancer cells. Through the detection of CIRI2, a total of 69,501 circRNAs were identified (Table S1). The length distribution of all identified circular RNAs revealed that over 80% of these circRNAs had lengths less than 2500 base pairs (bp) (Fig. 3A). In terms of circRNAs types, exonic circRNAs were the most predominant form, comprising 76.3% of the total, followed by exon-intronic circRNAs at 14.28% (Fig. 3B). To detect differentially expressed circRNAs, circRNA-pro utilized three software tools—Ciriquant, DEBKS, and CircTest, and detected 220, 1820, and 193 differentially expressed circRNAs, respectively. Notably, there was a large overlap in the circRNAs identified by these methods, with the largest overlap observed between CircTest and DEBKS, where 90% of the circRNAs detected by CircTest were also identified by DEBKS (Fig. 3C). Of the total 69,501 circRNAs identified, 40,671 were documented in CircBase and Circpedia database, indicating that 42% are novel discoveries (Fig. 3D). This reveals that numerous circRNAs remain to be discovered and investigated. According to the heat map, the differentially expressed circRNAs detected by circRNA-pro exhibit significant expression differences between breast cancer and healthy samples (Fig. 3E). To further investigate the functional implications of these circRNAs, we performed Gene Ontology (GO) enrichment analysis. Interestingly, the results revealed a significant enrichment in the immunoglobulin complex among the differentially expressed circRNAs (Fig. S1), suggesting a potential role for these circRNAs in the immune response associated with breast cancer. To gain a more comprehensive understanding of the genomic distribution and frequency of the differentially expressed circRNAs, we utilized the circRNA-pro to generate a circos plot (Fig. 3F). This visualization tool provides a detailed representation of the chromosomal locations and relative abundance of the circRNAs identified by various analytical tools, including Ciriquant, DEBKS, and CircTest.

**Fig. 3.**
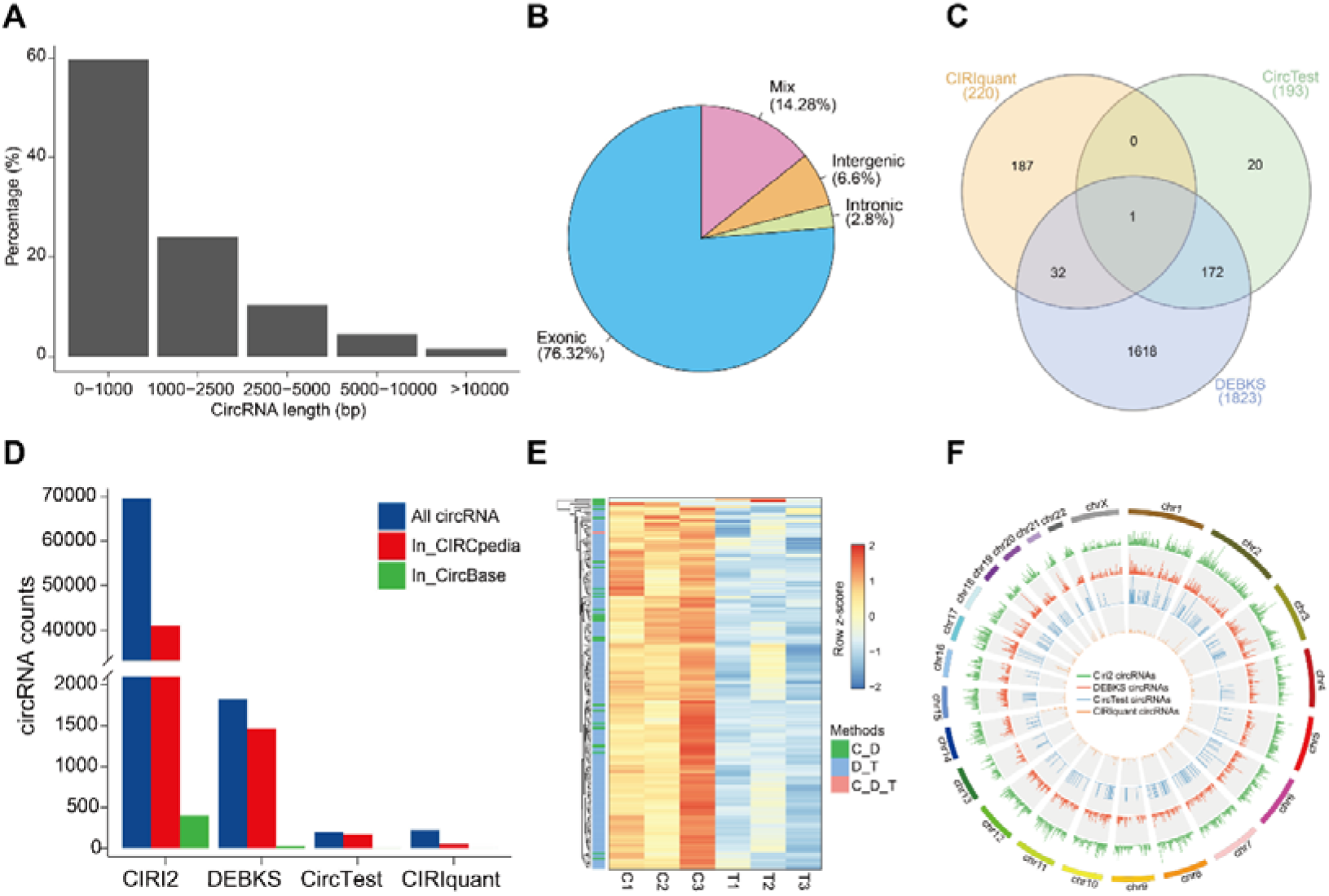
(A) Length distribution of circRNAs as identified by CIRI2. (B) Percentage of various circRNAs types. (C) Comparison of circRNAs detected by different software programs (Ciriquant, DEBKS, CircTest). (D) Comparative analysis of circRNAs identified by various methods against entries in circBase and CIRCpedia databases. (E) Differential expression of circRNAs in cancer vs. healthy samples, as detected by CircTest (T), DEBKS (D), and Ciriquant (C). (F) Chromosomal distribution and frequency of circRNAs detected by Ciriquant, DEBKS, and CircTest.

To identify circRNAs with translational capability, we aligned Ribo-seq reads to the sequence flanking circRNAs back-splice junction sites, initially screening out 943 circRNAs with potential for translation. Among these, 538 circRNAs had support from three or more Ribo-seq reads (Fig. 4A). To ascertain their translational potential, we cross-validated these 943 circRNAs against the riboCIRC and TransCirc databases, finding that 22 and 64 circRNAs, respectively, were previously recorded and supported by Ribo-seq evidence. Notably, 14 circRNAs were present in both databases (Fig. 4B). Of these candidates, the majority (75.8%) originated from exonic regions, while a minority (2.67%) were from intronic regions (Fig. 4C). Gene Ontology (GO) analysis of these translatable circRNAs candidates indicated significant enrichment in pathways associated with cell cycle regulation, apoptosis, and cell proliferation and migration, as depicted in Figure 4D.

**Fig. 4.**
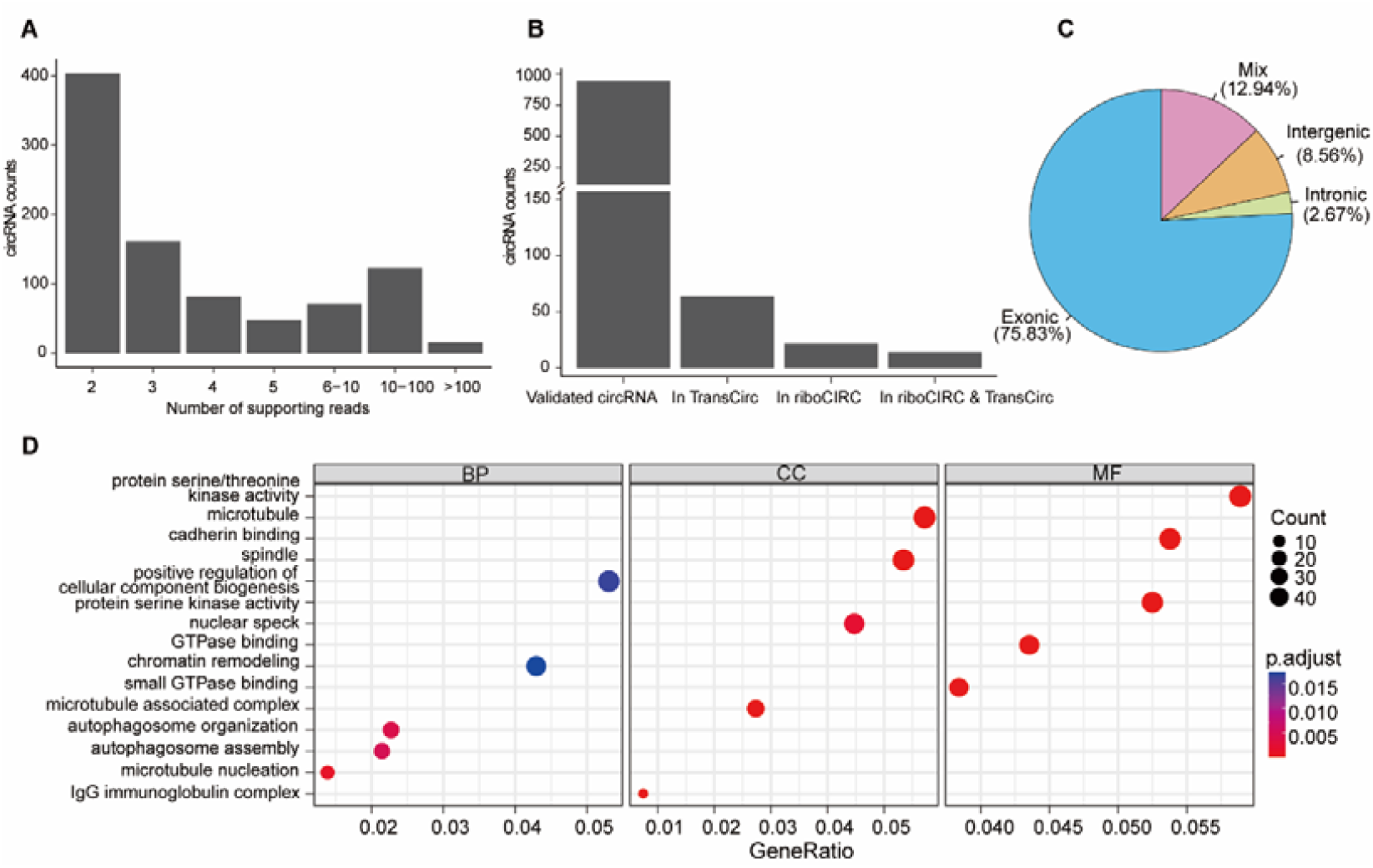
Analysis of potentially translatable circRNAs (A) Distribution of circRNAs supported by various numbers of Ribo-seq reads. (B) Comparison of translatable circRNAs identified by circRNA-pro with two translatable circRNAs databases. (C) Classification of translatable circRNAs candidates identified by circRNA-pro. (D) Gene Ontology (GO) enrichment analysis of translatable circRNAs identified by circRNA-pro.

### 2.4 The performance of circRNA-pro on the data of SARS-CoV-2 infected cells

SARS-CoV-2, a virus with severe threat for human health, has not been extensively investigated in terms of its impact on circRNAs in host cells [44, 45]. To explore this, circRNA-pro was utilized to detect differentially expressed circRNAs in SARS-CoV-2 infected peripheral blood mononuclear cells and Calu-3 cells compared to uninfected controls. In the peripheral mononuclear cells, CIRI2 detected a total of 18,302 circRNAs, among which 3640, 1, and 92 circRNAs were identified as differentially expressed by DEBKS, CIRIquant, and CircTest software, respectively (Table S2). Notably, there was a 95% overlap between the results of CircTest and DEBKS (Fig. 5A). In Calu-3 cells, out of 34,516 detected circRNAs, 2215, 7, and 47 circRNAs were identified as differentially expressed by DEBKS, CIRIquant, and CircTest, respectively (Table S3). Similar to peripheral blood mononuclear cells, a majority (87%) of CircTest results overlapped with DEBKS (Fig. 5B). Across the two cell types, 313 differentially expressed circRNAs were commonly detected by DEBKS, and 7 were commonly detected by CircTest (Fig. 5C).

**Fig. 5.**
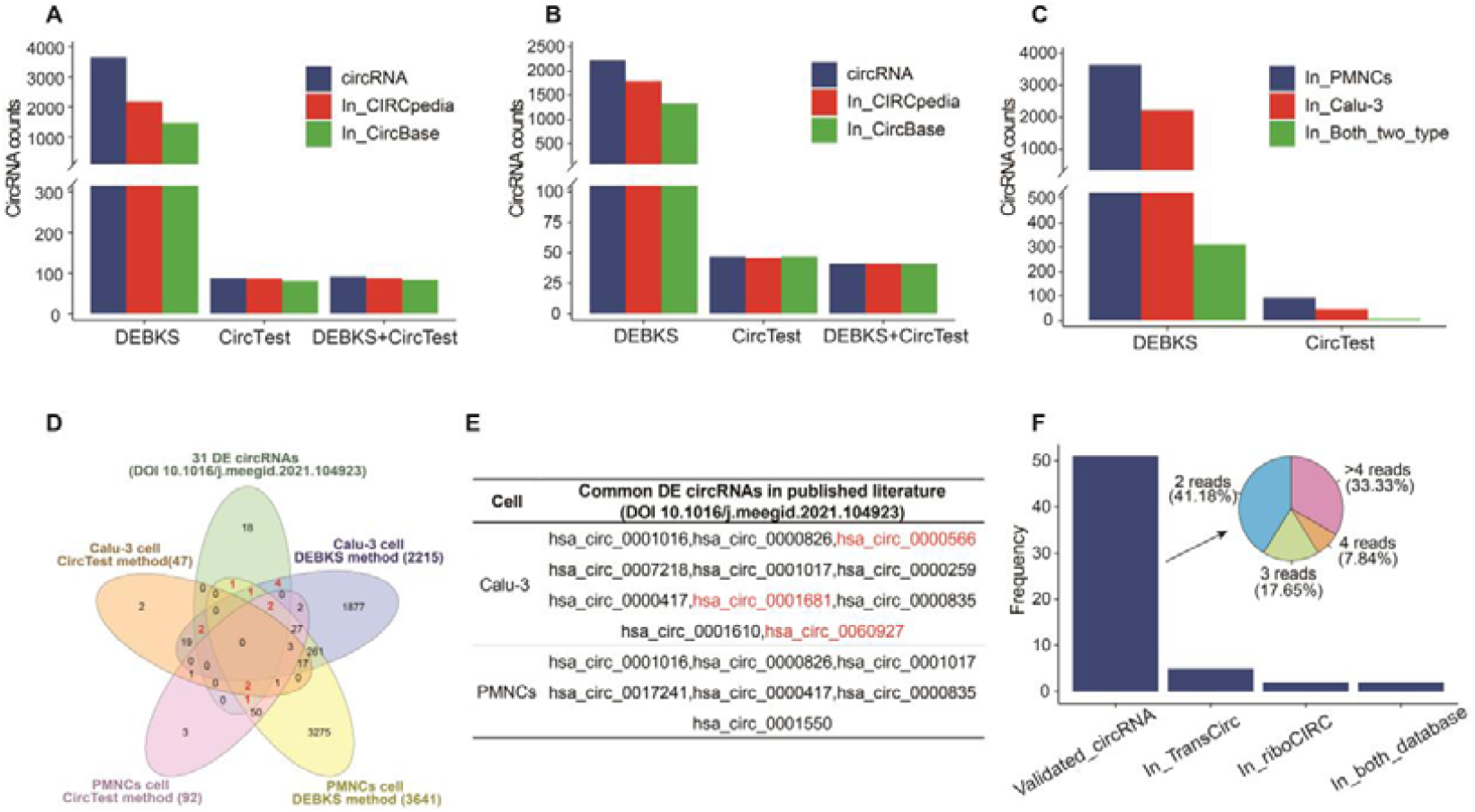
Differentially expressed circular RNAs and translatable circular RNAs in SARS-CoV-2-Infected cells. (A) Differentially expressed circRNAs in Peripheral Mononuclear Cells. (B) Differentially expressed circRNAs in Calu-3 Cells. (C) Differentially expressed circRNAs detected by circTest and DEBKS in different cell types. (E) Comparison of differentially expressed circRNAs in SARS-CoV-2 infected peripheral mononuclear cells and Calu-3 Cells with published literature. (F) Translatable circRNAs identified by circRNA-pro using Ribo-seq data, with comparison to two circRNA databases. A pie chart shows the percentage of translatable circRNAs with varying numbers of supporting Ribo-seq reads.

To further validate the accuracy, we compared the differentially expressed circRNAs identified in peripheral mononuclear cells and Calu-3 cells with those previously reported in human lung epithelial cells infected with SARS-CoV-2 [44]. Interestingly, 13 out of the 31 differentially expressed circRNAs identified in SARS-CoV-2 infected human lung epithelial cells [44] were also found to be differentially expressed in either peripheral blood mononuclear cells or Calu-3 cells (Figures 5D and 5E). Notably, among these 13 differentially expressed circRNAs, 3 circRNAs (hsa_crc_0000566, hsa_crc_0001681, hsa_crc_0060927) have been experimentally validated in human lung epithelial cells, further strengthening the reliability of the circRNA-pro detection results.

To identify potentially translatable circRNAs in SARS-CoV-2 infected cells, all identified circRNAs in Calu-3 cells were aligned with Ribo-seq data. A total of 51 circRNAs were identified with potential translational capability, among which 30 circRNAs (representing 58.8%) were supported by three or more Ribo-seq reads. Upon cross-validation with the Transcirc and Ribocirc databases, two public translatable circRNA databases, it was found that 5 of these circRNAs were documented in the Transcirc database, and 2 circRNAs (also recorded in the Transcirc database) were identified in the Ribocirc database (Fig. 5F). Notably, all these overlapping circRNAs were supported by Ribo-seq evidence in both databases.

## Discussion

In this study, we have developed a novel bioinformatics software specifically designed for the detection of differentially expressed circRNAs and potentially translatable circRNAs. CircRNA-pro can enhance the detection accuracy of these circRNAs through the integration of multiple analysis methods, while also being user-friendly. Firstly, circRNA-pro provides more reliable results for the detection of differentially expressed circRNAs by cross-validating the detection results from three distinct software algorithms. Secondly, the software incorporates an improved method for aligning Ribo-seq data with the sequences flanking circRNAs splice junctions to identify translatable circRNAs candidates. This includes removing sequences flanking circRNAs splicing sites that can be aligned to the reference genome or transcripts, as it is difficult to determine whether Ribo-seq sequences aligned to these overlapping regions originate from circRNAs or the reference genome. Furthermore, the software validates the potentially translatable circRNA candidates by comparing them against two commonly-used databases of translatable circRNAs. Moreover, circRNA-pro has been packaged into a Docker image, enabling easy distribution and execution across diverse computing platforms with a single command. This will facilitate the use by researchers with limited bioinformatics expertise and promote more discoveries and investigations into this emerging class of RNAs with important biological functions.

To assess the accuracy of circRNA-pro in detecting circRNAs, we applied it to the sequencing data from three distinct cell types: breast cancer cells, SARS-CoV-2 infected Peripheral Mononuclear Cells and Calu-3 cells. The software successfully identified a considerable number of circRNAs in each cell type—69,501 in breast cancer cells, 18,302 in PMNCs, and 34,516 in Calu-3 cells. Notably, a substantial fraction of these detected circRNAs were previously documented in well-established circRNAs databases (CircBase and Circpedia). The known circRNAs accounted for 58%, 61.2%, and 67.5% of the total circRNAs detected in each cell type, respectively. These results confirmed the accuracy of circRNA-pro in detecting circRNAs, while also indicating that there is still a large number of novel circRNAs to be discovered and investigated, as the known circRNAs documented in existing databases only accounted for 58-67.5% of the total circRNAs identified in each cell type.

After completing the detection of all circRNAs, circRNA-pro was then utilized to detect differentially expressed circRNAs from all the detected circRNAs using three different software tools, namely Ciriquant, DEBKS, and CircTest, in breast cancer sequencing data. The analysis results revealed significant discrepancies in the results obtained by each tool. DEBKS identified 1820 differentially expressed circRNAs, CircTest detected 193, while Ciriquant identified 220. Notably, there was a substantial overlap of 173 differentially expressed circRNAs between DEBKS and CircTest, but only minimal overlap with Ciriquant, which had 33 and 1 in common with DEBKS and CircTest, respectively. Further analysis of data from SARS-CoV-2-infected Peripheral Mononuclear Cells and Calu-3 Cells showed that Ciriquant detected an extremely limited number of differentially expressed circRNAs (1 and 7, respectively) with no overlap with either DEBKS or CircTest. In contrast, DEBKS detected 3641 and 2215 differentially expressed circRNAs, and CircTest detected 92 and 47, respectively, with a notable degree of overlap between DEBKS and CircTest (86 and 41, respectively). These results indicate that there are certain discrepancies among these software tools in detecting differentially expressed circRNAs. However, the overlap among them also highlights their consistency. The results from DEBKS and CircTest are relatively consistent with each other, whereas the results from Ciriquant significantly differ from the other two software tools. These observations not only demonstrate the divergence among software tools in the detection of differentially expressed circRNAs, but also highlights the importance of cross-validation and the necessity of employing multiple tools when confirming differentially expressed circRNAs, to ensure the accuracy and reliability of the findings.

To further validate the accuracy of circRNA-pro in detecting differentially expressed circRNAs, we compared differentially expressed circRNAs identified in SARS-CoV-2 infected PMNCs and Calu-3 cells with 31 differentially expressed circRNAs previously reported in SARS-CoV-2 infected human intestinal cells. The results showed that among the 31 differentially expressed circRNAs reported in the literature, circRNA-pro was able to detect 13 of them, with a 42% overlap rate. This indicates that circRNA-pro can effectively detect differentially expressed circRNAs, showing good consistency with the literature. More importantly, three differentially expressed circRNAs experimentally validated in the literature were also detected by circRNA-pro, further confirming its reliability in detecting differentially expressed circRNAs.

Furthermore, to evaluate the accuracy of circRNA-pro in identifying translatable circRNAs, we compared the putative translatable circRNAs identified by Ribo-seq data against two well-known databases of translatable circRNAs. In breast cancer and SARS-CoV-2 infected Calu-3 cells, we identified 943 and 72 potentially translatable circRNAs, respectively. Upon comparison with two widely-used translatable circRNAs databases, we found that 7.6% of the translatable circRNAs candidates from breast cancer cells and 9.8% from SARS-CoV-2 infected Calu-3 cells were previously recorded in these databases and supported by Ribo-seq evidence. These results not only validate the accuracy of circRNA-pro in detecting translatable circRNAs, as demonstrated by the overlap with the well-established RiboCirc and TransCirc databases, but also suggest that there may be a substantial number of novel potentially functional translatable circRNAs awaiting discovery and inclusion in current databases, which could have significant implications for our understanding of circRNA biology. It is also important to acknowledge that our current study relies on computational predictions and comparisons with existing databases, and future experimental validation is necessary to confirm the translational potential and biological functions of those identified circRNAs.

In summary, the development of circRNA-pro will significantly reduce the difficulty of circRNA analysis and improved the efficiency and accuracy of circRNA analysis results, providing critical support for a deeper exploration of circRNA function and regulatory mechanisms.

## Method

### RNA-seq and Ribo-seq datasets for the evaluation of circRNA-pro

In this study, we employed publicly available RNA-seq and Ribo-seq datasets to evaluate the performance of circRNA-pro. The RNA sequencing data and Ribo-seq data for breast cancer cells were obtained from BioProject PRJNA484546 [46] and PRJNA898352 [47], respectively. The RNA-seq data for PMNCs and Calu-3 cells infected with SARS-CoV-2 were sourced from BioProject PRJNA672981 [48] and PRJNA777558 [49], respectively. Additionally, the Ribo-seq data for SARS-CoV-2 infected Calu-3 cells were derived from BioProject PRJNA661467 [50].

### Implementation and installation of CircRNA-pro

CircRNA-pro is developed for the Linux platform, primarily using Python, and is available as a Docker image for ease of use. The Docker image is hosted on Docker Hub and can be accessed at https://hub.docker.com/repository/docker/songweidocker/circrna_pro. All experiments were performed on an Ubuntu Linux 18.04.3 server, powered by two Intel Xeon processors (each with 32 cores and a total of 64 threads) and equipped with 512 GB of RAM. Installing CircRNA-pro is straightforward by pulling the Docker image into the Linux system with the command: docker pull songweidocker/circrna_pro:v1. It is necessary to have Docker installed on the Linux system before installing CircRNA-pro. The installation instructions for Docker can be found at: https://docs.docker.com/engine/install/.

### Detection and visualization of differentially expressed circRNAs

To identify circRNAs, we initially employed FastQC software for quality control of sequencing reads, generating a quality control report to assess the read quality. Subsequently, Fastp [51] software was utilized for further preprocessing of the reads, including quality filtering and adapter trimming, to produce high-quality clean reads. We then employed CIRI2, a software specifically designed for circRNAs identification, to align these preprocessed clean reads to the reference genome. CircRNAs were identified by detecting reads with reverse splicing junctions, using the aligned reads data. Following circRNAs identification, we used a suite of tools including Ciriquant [37], CircTest [39] and DEBKS [38] to detect differentially expressed circRNAs between samples. The expression levels of differentially expressed circRNAs identified by these tools across various samples were visualized using the Pheatmap package. Furthermore, the eulerr R package was utilized to compare the common and unique differentially expressed circRNAs identified by each of these tools.

### Identification and verification of translatable circRNAs

To identify translatable circRNAs, we implemented a multi-strategy approach, combining Ribo-seq sequence alignment with comparison verification against translatable circRNAs databases. Ribo-seq reads were first aligned to the reference genome using Tophat2 [52]. The unmapped reads were retained for downstream analysis, while mapped reads were filtered out. We then constructed a pseudo circRNAs genome by concatenating 10 base pair (bp) sequences flanking circRNAs splicing sites. These concatenated sequences were aligned to the reference genome and transcripts using blastn [53] to remove sequences aligning to genomic or transcriptomic regions. The filtered Ribo-seq reads were subsequently aligned to the pseudo genome of circRNAs. CircRNAs supported by at least two aligned Ribo-seq reads were preliminarily identified as translatable. As further validation, the putative translatable circRNAs candidates were compared to two widely recognized translatable circRNAs databases, namely RiboCirc and TransCirc.

### Comparing circRNAs with four circRNAs databases

After identifying circRNAs candidates using circRNA-pro, we conducted comparative analyses against four well-known circRNAs databases: CircBase [54], Circpedia [55], Ribocirc [40], and TransCirc [56]. To ensure the most up-to-date results, circRNA-pro allows for flexible comparison with either locally downloaded database or online databases. To ensure compatibility with the genomic version used by circRNA-pro, the coordinates of circRNAs in the database entries were adjusted to match the genomic version employed by the circRNA-pro using the liftOver [57] tool. CircRNAs identified by circRNA-pro were considered identical to those in the databases if their genomic coordinates varied by less than 10 bp.

### The visualization of CircRNA-pro results

To visualize the results obtained from circRNA-pro, we employed the VennDiagram [58] R package to create Venn diagrams illustrating the intersections between differentially expressed circRNAs, translatable circRNAs, and circRNAs from relevant databases. Furthermore, a circos plot was constructed using the circos [59] R package to provide a comprehensive depiction of the circRNAs landscape across the genome. This plot clearly presents the chromosomal locations and relative abundance of the circRNAs identified by various analytical tools, including Ciriquant, DEBKS, and CircTest.

## Supporting information

Supplementary_Figure_1_GO enrichment_analysis_of_differentially_expressed_circRNAs_in_breast_cancer_cells

Supplementary_Table_1_Differentially_Expressed_circRNAs_and_Translatable_circRNAs_in_Breast_Cells

Supplementary_Table_2_Differentially_Expressed_circRNAs_and_Translatable_circRNAs_in_SARS-CoV-2_Infected_peripheral_mononuclear_cells

Supplementary_Table_3_Differentially_Expressed_circRNAs_and_Translatable_circRNAs_in_SARS-CoV-2_Infected_Calu-3_Cells

## Author contributions

Conceptualization, W.S., H.J. and W.P.; the development of CircRNA-Pro program and data analysis, W.S., L.Y. and T.Y.; writing—draft preparation, W.S., Y.W. and Y.Y.; writing—review and editing, W.P., H.Z., D.S. and H.J.; supervision, H.J., W.P.. All authors have read and agreed to the published version of the manuscript.

