## Supplementary figures and images for "CircRNA-Pro: A Novel Toolkit for High-Precision Detection of Differentially Expressed Circular RNAs and Translatable Circular RNAs"

### Supplementary_Figure_1_GO enrichment_analysis_of_differentially_expressed_circRNAs_in_breast_cancer_cells

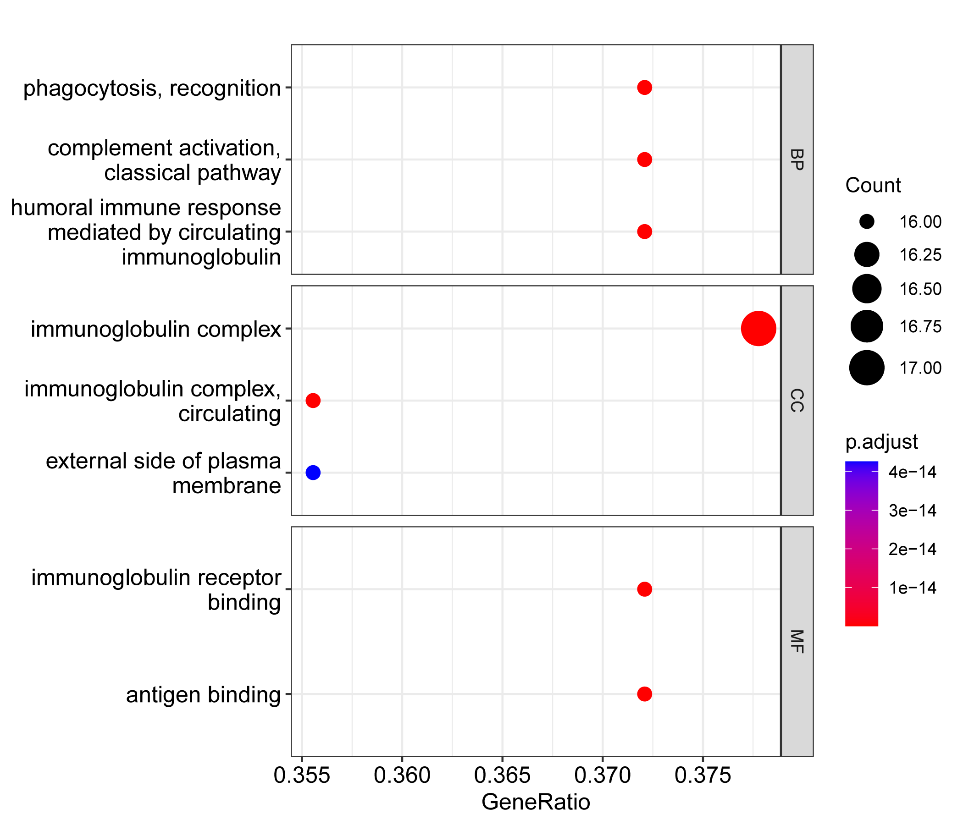

Fig. S1 GO enrichment analysis of differentially expressed circRNAs in breast cancer cells
